# The characteristics of high-dark-diversity habitats derived from lidar

**DOI:** 10.1101/2022.05.05.490326

**Authors:** Jesper Erenskjold Moeslund, Kevin Kuhlmann Clausen, Lars Dalby, Camilla Fløjgaard, Meelis Pärtel, Norbert Pfeifer, Markus Hollaus, Ane Kirstine Brunbjerg

## Abstract

A key aspect of nature conservation is knowledge of which aspects of nature to conserve or restore to favor the characteristic diversity of plants in a given area. Here, we used a large plant dataset with > 40.000 plots combined with airborne laser scanning (lidar) data to reveal the local characteristics of habitats having a high plant dark diversity – i.e., absence of suitable species – at national extent (> 43.000 km^2^). Such habitats have potential for reaching high realized diversity levels and hence are important in a conservation context. We calculated 10 different lidar based metrics (both terrain and vegetation structure) and combined these with 7 different field-based measures (soil chemistry and species indicators). We then used Integrated Nested Laplace Approximation for modelling plant dark diversity across 33 North European habitat types (open landscapes and forests) selected by the European communities to be important. In open habitat types high-dark-diversity habitats had relatively low pH, high nitrogen content, tall homogenous vegetation and overall relatively homogenous terrains (high terrain openness) although with a rather high degree of local microtopographical variations. High-dark-diversity habitats in forests had relatively tall vegetation, few natural-forest indicators, low potential solar radiation input and a low cover of small woody plants. Our results highlight important vegetation, terrain and soil related factors that managers and policymakers should be aware of in conservation and restoration projects to ensure a natural plant diversity, for example low nutrient loads, natural microtopography and open forests with old-growth elements such as dead wood and rot attacks.

## Introduction

A key aspect of nature conservation is knowledge of which aspects of nature to conserve or restore to favor the characteristic diversity of plants in a given area (Hanson et al. 2020; Penone et al. 2019). Studying the diversity of species that are absent, despite suitable conditions – the dark diversity (Pärtel et al. 2011) – could possibly aid in pointing out such aspects (Lewis et al. 2017). Here, we use a massive national plant dataset combined with airborne laser scanning (lidar) data to reveal the characteristics of habitats having a high dark diversity of plants, and we discuss how this knowledge can augment practitioners’ ability to efficiently conserve important biotopes.

Lidar has ample potential for characterizing terrain and vegetation structural features of a habitat and to some degree also the abiotic environment (Moeslund et al. 2019). Today, the details from lidar data are so good that vegetation structure even in non-woody habitats with low vegetation can often be retrieved. For this reason, lidar is increasingly used in studies of local-scale ecology and biodiversity (de Vries et al. 2021; Moeslund et al. 2019; Mäyrä et al. 2021), although it has never been used for studying plant dark diversity (but see Valdez et al., 2021).

Most studies of biodiversity use a measure based on observed diversity – for example species richness – as the sole measure of biodiversity. This approach ignores the dark diversity; the species that could thrive in a site but is absent currently, and consequently it also fails to convey information on why these species are missing. Some natural areas may harbor a relatively high diversity and yet at the same time have a high dark diversity (see for example Morel et al., 2022), but such knowledge is not revealed in studies focusing only on present diversity. This also makes traditional observed-diversity studies unsuitable for cross-habitat comparisons of the biodiversity as different habitats have different species pools (Zobel 2016).

The concept of dark diversity is ideal to study the relationship between environmental properties and potentially missing species in a given habitat in order to learn how to improve its status towards more natural conditions. The field of dark diversity is still rather new but several studies have used it to study characteristics of either habitats having a high dark diversity or species having a high tendency for belonging to the dark diversity (Moeslund et al. 2017; Penone et al. 2019; Riibak et al. 2015; Wen et al. 2020). These studies indicated that for plants, open-landscape habitats having low mycorrhizal diversity and high nutrient and stress levels are often associated with a relatively high dark diversity (Moeslund et al. 2017; Riibak et al. 2015). Notably, the study by Penone et al. (2019) highlighted the possible importance of vegetation structures for the dark diversity in forests. However, most of these studies covered only a few habitat types, or included only a relatively small number of study sites or species, preventing a broader application of results in conservation management.

Our study is the first to use lidar for studying plants’ dark diversity, and we present results that are broadly relevant across temperate European habitats – e.g., from wet to dry, open to forested, acidic to calcareous. We used vegetation data from approximately 43.000 vegetation plots distributed over approx. 42.000 km^2^ combined with field- and lidar-based measures to examine the characteristics of habitats having a high dark diversity of plants. More specifically, we addressed the following questions: (1) What characterizes habitats with a high plant dark diversity in temperate northern Europe, and (2) which factors are the most important in this regard? (3) Are field-based measurements needed in modelling local plant dark diversity or does remote sensing (lidar-based measures) alone perform just as well?

## Methods

All statistical analysis and data handling described below were conducted in the R environment (Team 2020) unless otherwise stated.

### Plant data

To enable estimation of plant dark diversity, we used presence-absence data from the habitats monitoring program for Denmark (NOVANA, novana.au.dk). Through this program, the presence of all plant species in 5-m (open landscape) or 15-m (forest) circular vegetation plots is recorded for > 50.000 plots covering all Danish habitat types listed in Annex I of the European Union’s Habitats Directive (Council of the European Communities 1992). In November 2018, we extracted data for 53.174 plots for the monitoring period 2004 – 2018 (naturdata.miljoeportal.dk). We removed lakes and streams to focus on terrestrial habitats. We removed all records of collective species (e.g., *Taraxacum officinale* coll. and *Rubus fruticosus* coll.). Subspecies and varieties were lumped to the species level. We also removed all records of neophytes according to Buchwald et al. (2013) and plots having less than five species. Then, we aggregated plot data from different years, assuming that the vegetation has not changed notably over the period. Approximately 28 % of the plots have been visited only once, 40.5 % two to four times, 21 % five to six times, 10 % seven to nine times and 0.5 % ten to twelve times. Finally, we added 448 vegetation plots (nature or semi-nature) similar to the above from a national scientific biodiversity project; Biowide (Brunbjerg et al. 2019). Hereafter, the dataset consisted of 43,703 plots with 1,002,663 records of 1,089 species.

### Dark diversity

To calculate the dark diversity for each plot, we first calculated the site-specific species pool using Beals’ index (Beals 1984), as recommended by Lewis et al. (2015), using the “beals” function in the “vegan” package (Oksanen et al. 2017). Beals’ index represents the probability that a particular species will occur based on the assemblage of co-occurring species (Beals 1984; Münzbergová and Herben 2004). The site-specific species pool was estimated as follows. The threshold for including a particular species in the sitespecific species pool was the 5^th^ percentile of the Beals’ index value for this species following Gijbels et al. (2012) and Ronk et al. (2015). Before calculating these thresholds, we identified the minimum Beals’ index among plots with occurrence of the species in question and disregarded plots having values below this minimum. Subsequently, we calculated the dark diversity as the absent part of the site-specific species pool. Since species pools vary across environmental gradients, we divided the plot dark diversity by the site-specific species pool to allow cross-plot analysis. We refer to this measure as the *species-pool-adjusted dark diversity* in the following.

### Environmental data

To enable a proper habitat characterization, we used both field-measured and lidar-based measures in our statistical analysis. For some vegetation plots, several environmental factors were recorded in the field as part of the monitoring program, for example soil pH and soil nitrogen content (Table 1). Field-data collection procedures are described in the Supporting Information.

**Table 1.**
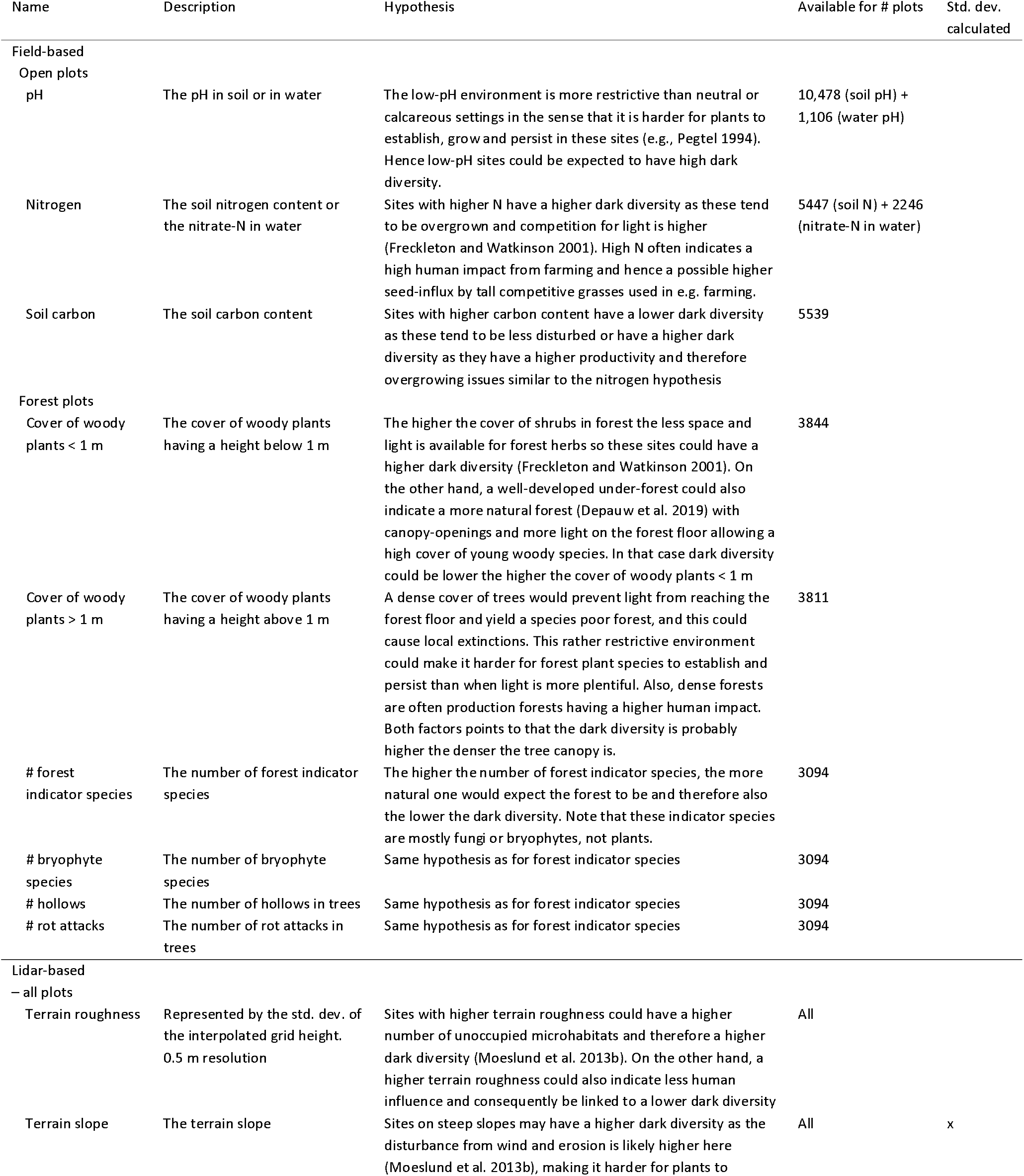
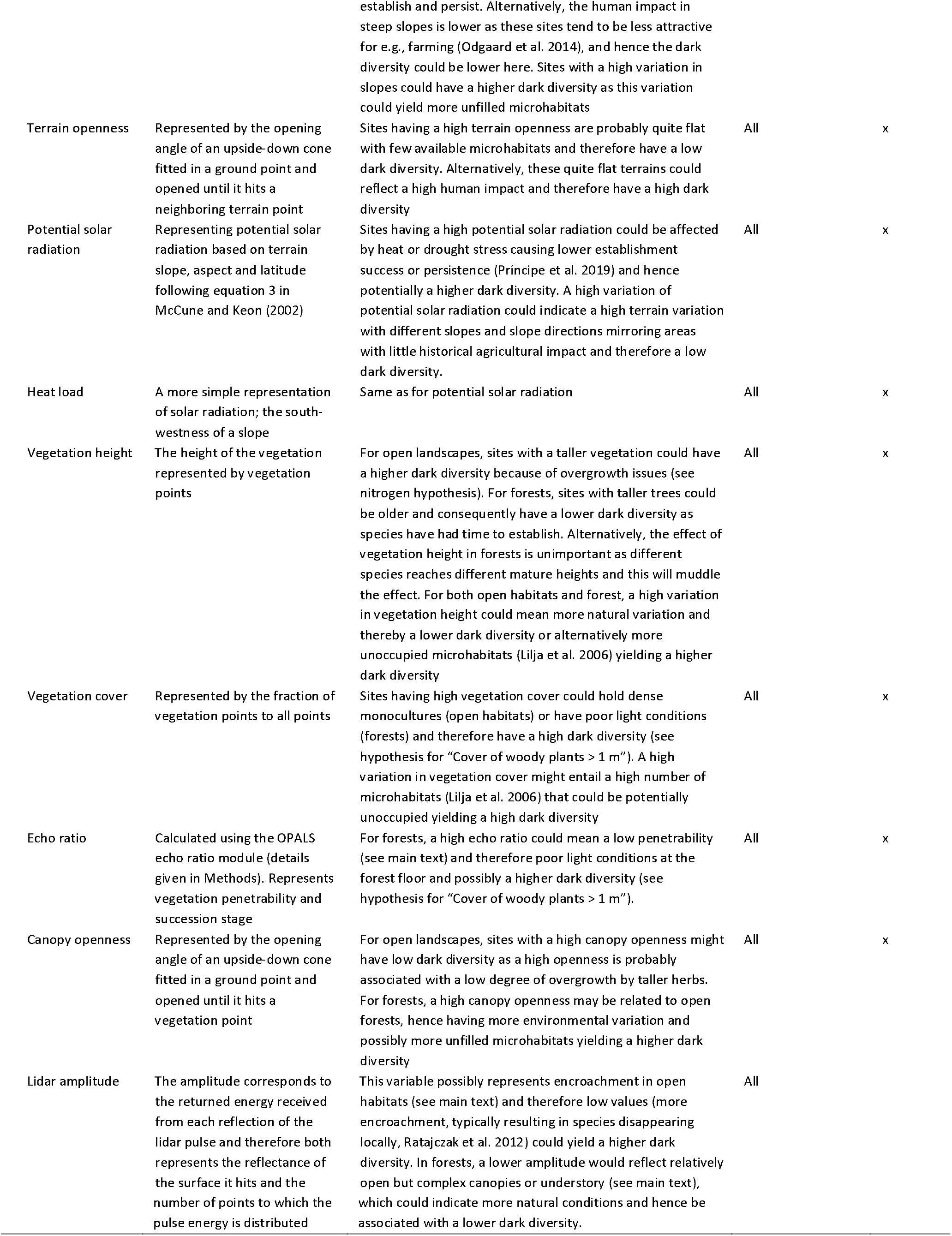
Overview of all explanatory variables used in this study, with a brief description, hypotheses, availability and indication of whether or not their standard deviation was also used in modelling (x).

For all vegetation plots, we calculated a number of lidar-based measures reflecting both the abiotic environment and terrain and vegetation structures. These were all based on the newest lidar point cloud for Denmark, recorded in springs and autumns of 2014 and 2015 (freely available at datafordeler.dk). The point cloud has a density of 4–5 points/m^2^ and was recorded by Riegl LMS-680i scanners (1550 nm) operating in a parallel line scan pattern, onboard fixed-wing airplanes (altitude: 680 m above ground level, speed over ground: 240 km/h). For all calculations, we used the data provider classification of points into ground, building and vegetation classes. We calculated all measures at 1.5 m grid spacing (except for terrain roughness, which was at 0.5 m) and their means and standard deviations within 30 m radius circles centered in each vegetation plot. For all lidar processing and calculations, we used the OPALS tools version 2.3.1 (Pfeifer et al. 2014) in a Python 2.7 environment. For subsequent geospatial calculations, we used the GDAL 2.2.2, rasterio 1.0.13 and geopandas 0.5.0 packages.

### Vegetation-related measures

To represent succession and – to some degree – moisture balance in both vegetation and soil, we used the *amplitude* of each echo representing a point in the lidar point cloud. This amplitude is high if the reflecting surface is flat and has a high reflectivity. It is low when the light energy is distributed between several returns for example in tree canopies. The 1550 nm wavelength is sensitive to leaf water content (Junttila et al. 2018) and soil moisture (Zlinszky et al. 2014). To account for variation due to different aircraft types and recording dates, we used the residuals of a Generalized Linear Model (GLM) having amplitude as response and flight month and airplane as explanatory variables. We refer to this as *corrected amplitude* in the following. We did not have reference data enabling a full calibration of this measure.

To represent *vegetation height,* the local terrain model was subtracted from the local surface model (detailed in the following). The terrain model (DTM) calculation details are given in the section “Terrain-structure measures”. The surface model was calculated using the “DSM” module in OPALS and all non-building points.

To reflect the vegetation penetrability and succession, we calculated the *echo ratio* (Höfle et al. 2012). Echo ratio is high where the surface is impenetrable and relatively smooth and lower where the surface is uneven or penetrable. For this calculation we estimated normals for each point using the “Normals” module in Opals with a robust plane fit based on the 12 nearest neighboring points. Echo ratio calculations for each terrain and vegetation point used the slope adaptive method implemented in the “Echo Ratio” module of Opals (search radius: 1.5 m).

To estimate light conditions, we calculated the *canopy openness* for all ground points. This is the same as terrain openness (see details below) but considering both ground and vegetation points in the process. Hence, canopy openness represents the actual blocking of the sky view by the canopy around each ground point. Canopy openness is high for ground points inside canopy gaps, and low for ground points beneath a closed canopy.

Lastly, as an estimate of *vegetation cover, we* calculated the fraction of vegetation points to all non-building points. This measure is high if the vegetation cover is high and dense, and low for areas with little vegetation.

### Terrain-structure measures

To enable the calculation of terrain-related measures, we calculated a digital terrain model (DTM) for each plot representing the elevation above sea level. To do this, we used the “DTM” module of Opals using 8 neighboring points and a search radius of 6 m. To represent key features of the local terrain (e.g., soil moisture or heat balance, Moeslund et al., 2013a), we calculated *terrain slope* and *terrain aspect* using the “GridFeature” module of Opals taking the DTM as input and a kernel size of 3×3 pixels.

To reflect local heat input, we calculated the *heat load index* following McCune and Keon (2002). This index reaches maximum values on southwest-facing slopes and zero on northeast-facing slopes. We also calculated the *potential solar irradiation* (over a year) following equation 3 in McCune and Keon (2002).

To estimate micro-scale *terrain roughness, we* calculated the standard deviation of the interpolated grid height (also termed *SigmaZ).* The Opals “DTM” module outputs this measure as a bi-product. Unlike our other lidar measures, *terrain roughness* was calculated at 0.5 × 0.5 m resolution mirroring micro-scale terrain variations.

To represent plot-scale terrain heterogeneity, we calculated the *terrain openness* (Doneus 2013). Terrain openness is defined as the angle of a cone turned upside down – with its tip restrained to the point of interest – when it touches the points closest to the surface normal vector. To calculate this, we used the “PointStats” module of Opals requesting “positive openness” based on ground points only and a search radius of 5 meters. This measure is high in flat (relative to the scale at which it is calculated) areas and low in heterogeneous terrains.

Finally, to test the importance of variations, we calculated the standard deviation for lidar measures for which we believed it made ecological sense (Table 1).

### Data preparation

Initially, all explanatory variables were visually checked for spatial patterns and outliers. Based on this, we removed data from the Island of Bornholm which is an outlier in terms of environment, soil type and geography. We also removed data from *Cladium mariscus* fens (canopy openness behaves strangely here, 320 records), and 136 records with outliers in lidar amplitude (values above 650) or its standard deviation (values above 350) as well as 22 records with outliers in terrain roughness (values above 0.08). Figure 1 shows the plots used in this study.

**Figure 1.**
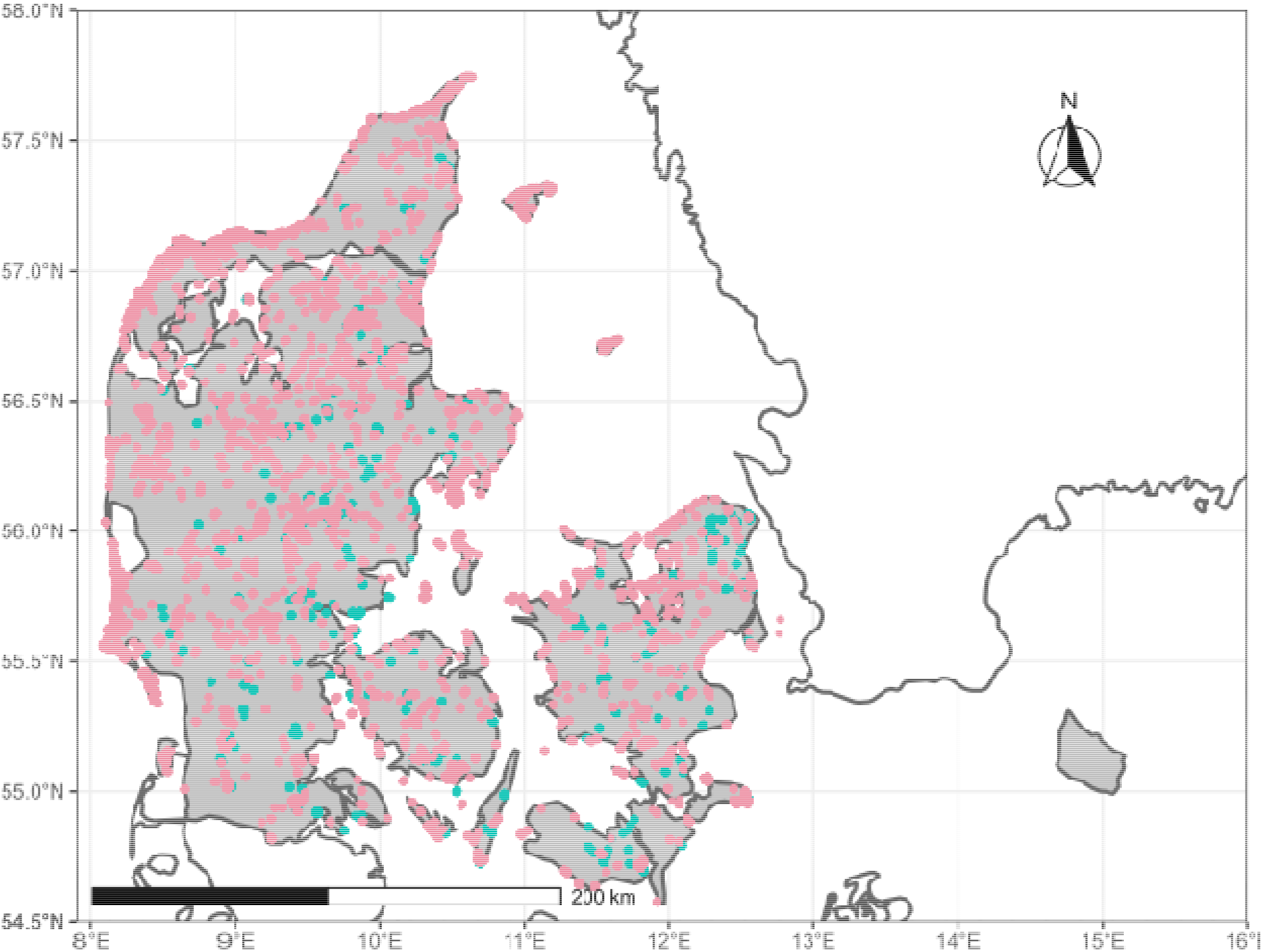
Overview of the vegetation plots included in this study. Red dots represent open landscape plots whereas green dots are forest plots.

Several of the measured variables were only available for a subset of all plots (Table 1) and their respective availability was not always overlapping. Therefore, we identified the set of measured variables with highest combined availability and best complementing the lidar variables (i.e., soil- and species-related measures as these are not well represented by lidar-derived measures). For forests, the best such combination was (1) coverage of woody plants under 1 m and (2) over 1 m, (3) number of forest indicator species, (4) number of bryophyte species, (5) number of living trees with hollows (holes in the bark with underlying rot having a depth of > 5 cm), and (6) number of living trees with rot attacks (areas > 100 cm^2^ with bark falling off and clear signs of decomposition). These were available for 3.094 forest plots. For the open landscapes the best combination was: (1) soil or water pH, (2) soil carbon content and (3) soil nitrogen or water nitrate-N content. These were available for 6.486 open-landscape plots.

To prevent multicollinearity issues, we ran a Variance Inflation Factor (VIF) analysis and left out explanatory variables with a VIF value above 5 (Menard 2001, Table S1 in Supporting Information).

### Statistical analysis

Statistical analysis was divided into two parts: open landscape and forests (Figs. S3 and S4 in Supporting Information). From prior experience, we knew that modeling all plots together would give strong importance to lidar variables separating the plots into open and forested vegetation and thereby obscure the importance of other variables. Because pH and nitrogen were measured differently over the years in some habitats, the open-landscape analysis was further subdivided into primarily dry habitats (pH and nitrogen measured in soil) and wet habitats (pH and nitrogen measured in water samples). We used only explanatory variables for which we had hypotheses regarding their relationships to dark diversity (Table 1). Firstly, we used GLMs with Gaussian error distribution. We used the regional pool adjusted dark diversity (see section “Dark diversity”) as response variable and the field-measured and lidar-derived variables as explanatory variables. To strive for normal distribution of explanatory variables we transformed those where this made obvious distributional improvements (visual examination of histograms, see Table S1 in Supporting Information for details). To enable comparison of coefficient estimates, explanatory variables were standardised (mean = 0, standard deviation = 1). To check for possible non-linear relationships, we tested the effect of squared terms for explanatory variables where we had a non-linear hypothesis (e.g. hump-shaped). We used Akaikes Information Criterion (AIC, Akaike 1974) to do this. If AIC of the new model was > 2 lower than the original (following Burnham and Anderson 2002), we kept the squared term for further model selection. Subsequently, we ran a backwards model selection procedure based on AIC always leaving out the explanatory variable that caused AIC to drop the most, but finally also leaving out variables (still one at a time) whose absence from the model did not cause AIC to increase by more than 2, following Burnham and Anderson (2002).

By calculating Moran’s I (using the function “Moran.i” of the “ape” package, Paradis and Schliep 2018) of the model residuals, we detected significant spatial autocorrelation for all models. We used Integrated Nested Laplace Approximation (INLA, Rue et al. 2009) models as described below to account for this (“R-INLA” package, www.r-inla.org), using the stochastic partial differential equation (SPDE) approach (Lindgren et al. 2011). Initially, we ran non-spatial INLA models corresponding to our GLM models. Then, we followed the principles described in Blangiardo et al. (2013) and outlined in the following. For all INLA models, we first created a triangulated mesh using “inla.mesh.2d” with a max edge of 50 km, a cut-off of 2 km and a boundary reflecting the terrestrial area of Denmark. This represented the spatial structure in our data.

Then we constructed the observation matrix A using “inla.spde.make.A” with the mesh and the locations of observations. Subsequently, we created a Matern SPDE model object with “inla.spde2.pcmatern” taking the mesh as input, and we created model index vectors using “inla.spde.make.index”. We now stacked the above into an “inla.stack” in preparation for INLA modelling. To test the degree to which accounting for spatial autocorrelation improved our models, we first added ID random effects by adding an ID column with integers from 0 to the number of records in the modelled dataset and treated these as a random effect in the model (model = “iid”). Subsequently, we added SPDE random effects (model = SPDE) on top of the ID random effects. We compared these two approaches by the deviance information criterion (DIC). Marginal posterior distributions were summarised by 95% Bayesian credible intervals (BCI) corresponding to the 0.025 and 0.975 quantiles of the posterior distribution (Zuur et al. 2017).

## Results

INLA models with SPDE random effects strongly improved model fit (DIC) in all cases (Table 2) with the main autocorrelation found within approx. 2–5 km for most open and forest habitats and 7–20 km for the wet open habitats (Figs. S5 and S6). Hence, we only report results from these models.

**Table 2.** Characteristics of the Integrated Nested Laplace Approximation models used to evaluate the relationships between local plant dark diversity and field- and lidar-based measures.

| Model | Sample size (n) | DIC (original) | DIC (w/ spatial random effect) |
| --- | --- | --- | --- |
| Open landscape |  |  |  |
| Dry |  |  |  |
| Measured + lidar | 5,380 | -14,928.56 | -19,293.10 |
| Lidar only | 5,380 | -13,779.00 | -19,093.40 |
| Wet |  |  |  |
| Measured + lidar | 1106 | -3,519.92 | -4,281.11 |
| Lidar only | 1106 | -3,282.29 | -4,137.94 |
| Lidar only, all | 38,244 | -108,322.21 | -138,583.27 |
| Forests |  |  |  |
| Measured + lidar | 3,094 | -10,949.65 | -13,568.95 |
| Lidar only, reduced | 3,094 | -10,480.48 | -13,640.10 |
| Lidar only, all | 3,844 | -12,618.35 | -16,263.02 |

Generally, for open habitats high-dark-diversity sites were characterized by having relatively low pH, tall homogenous vegetation and overall relatively homogenous terrains (high openness) although with a rather high degree of local variations (terrain roughness, Figs. 2a and 3a). In the dry open habitats, the plant dark diversity was also higher in sites with relatively high soil nitrogen content, in sites with a homogenous vegetation layer (low canopy openness variation) and in places with homogenous terrain openness (Fig. 2a). In wet open habitats, sites with little variation in solar radiation and sites with rather heterogeneous terrain openness had a higher plant dark diversity than others (Fig. 2a).

**Figure 2.**
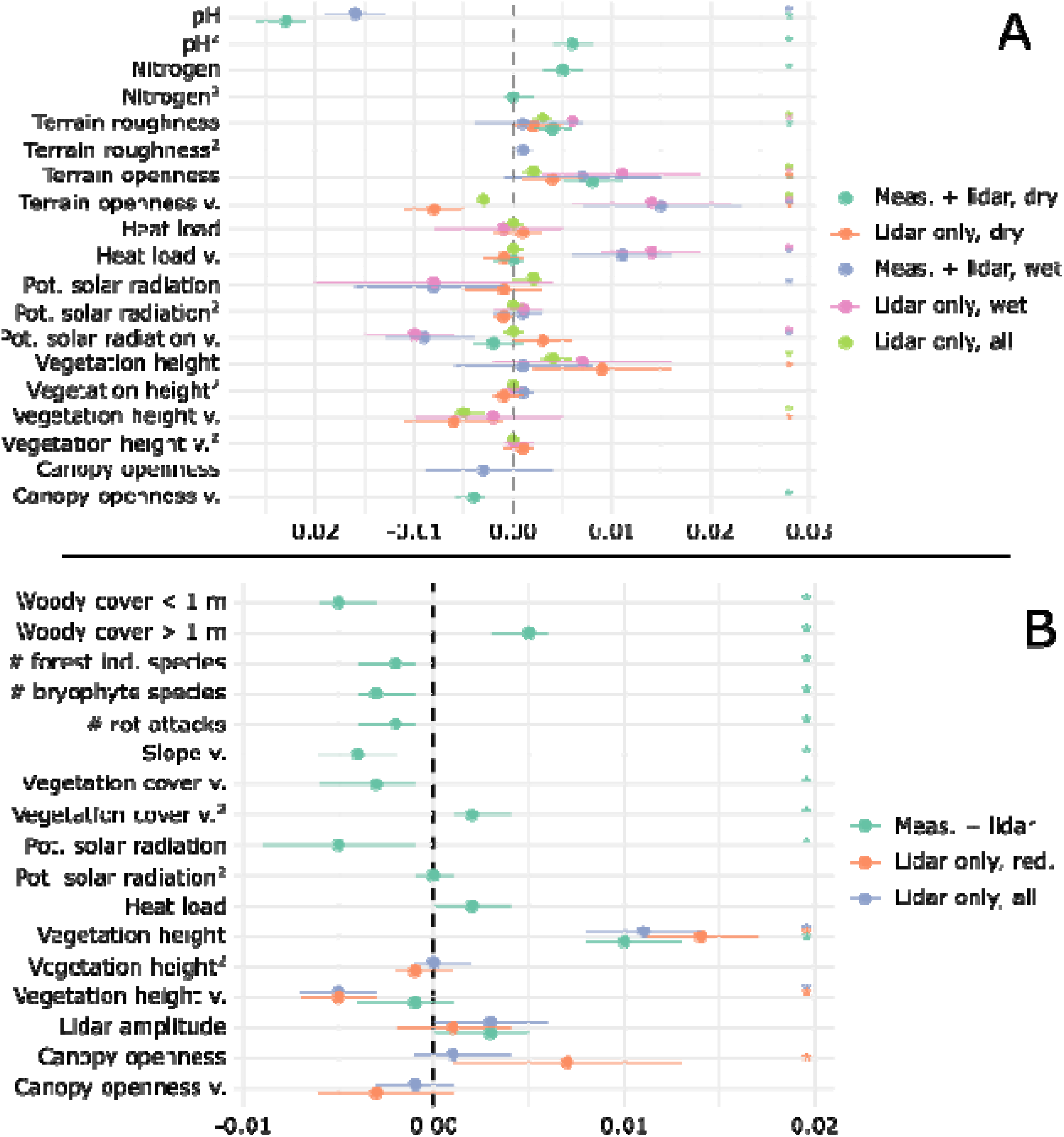
Standardised coefficient estimates from the Integrated Nested Laplace Approximation modelling of regional pool adjusted plant dark diversity in open (A) and forest landscapes (B). Asterisks indicate that the given variable was statistically significant (i.e., its confidence interval does not overlap with zero) in the model denoted by its color. Abbreviations: v. = variation, Pot. = potential, # = number of, ind. = indicator, red. = reduced, Meas. = measured

**Figure 3.**
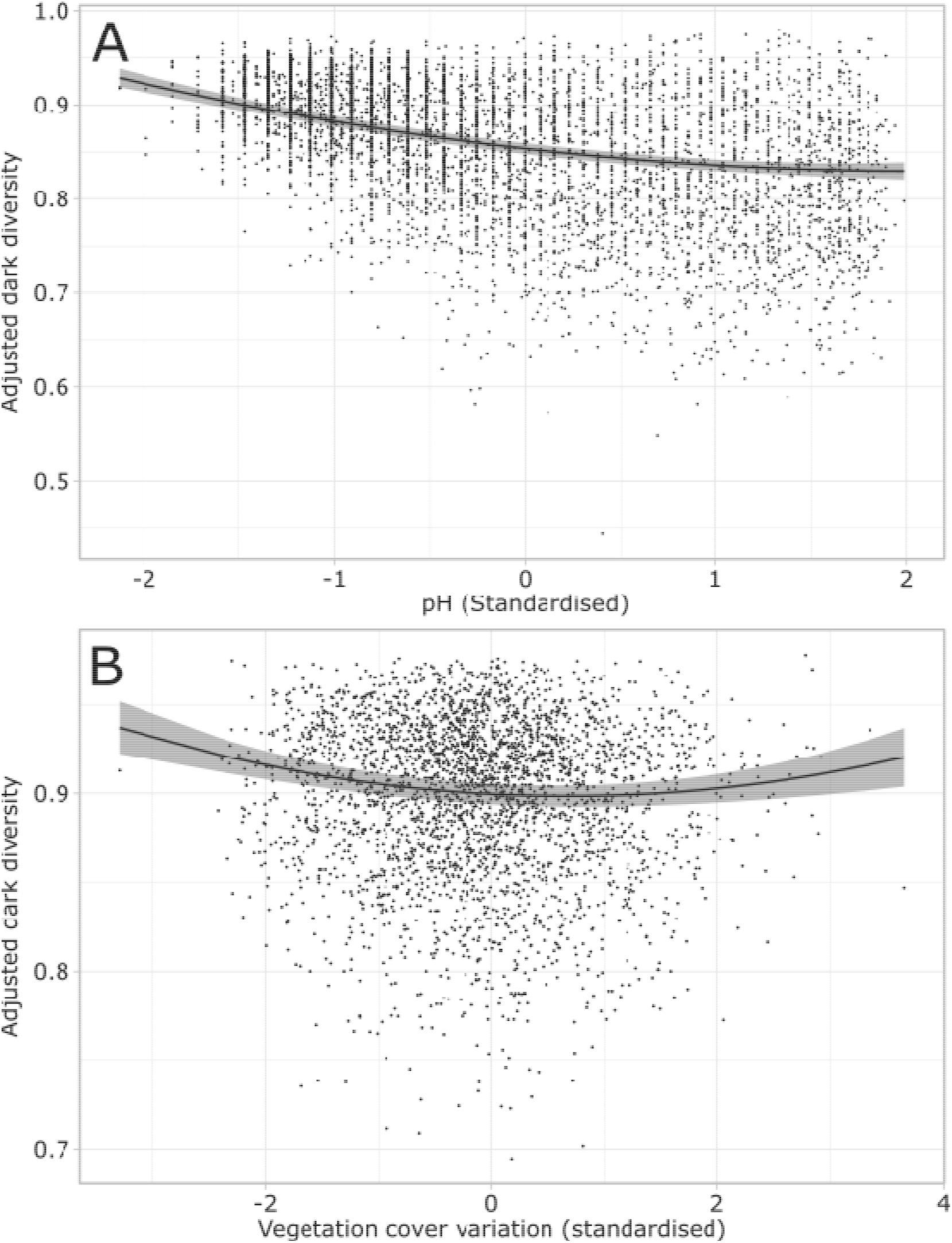
The effect of (A) pH in the model for open dry habitats and (B) of the variation (standard deviation) in vegetation cover on in the model for forest habitats when all other factors in the model were held constant at their mean values. Both the predicted values (black line) and the 0.025 and 0.975 quantile values (gray surface) are shown.

In forests, high-dark-diversity sites were characterized by relatively tall vegetation and fewer natural-forest indicators (number of forest indicator species, bryophyte species and rot attacks, Fig. 2b). Also, forest sites with a high dark diversity had a low solar radiation input and they had a low cover of small woody plants (< 1 m, Fig. 2b). The variations in vegetation cover showed a non-linear relationship to plant dark diversity in forests, with lowest dark diversity at intermediate variations and higher towards the extremes (Figs. 2b and 3b).

The most important factor affecting local plant dark diversity in open landscapes was soil pH, while in forests vegetation height was the most important (Fig. 2). Vegetation height and variations therein were consistently important for local plant dark diversity both in open landscapes and in forests (Fig. 2).

Models consistently performed better with measured variables included than if lidar based variables were the only explanatory factors (Table 2).

The detailed modelling results including sample size, DIC values and posterior means and 0.025 and 0.975 quantiles are given in Table 2 and Figure 2.

## Discussion

### High-dark-diversity habitat characteristics

#### Open habitats

Our analyses showed that open habitats with a relatively low pH also tend to have a higher plant dark diversity. It is well-known that high-pH soils in Europe are species rich because of the high species pool associated with these soils (Pärtel 2002). In addition to this, we now show that plant dark diversity is relatively low in these high-pH soils, suggesting that there are two mechanisms causing high-pH habitats to be species rich. The high dark diversity on low-pH soils could be explained by low-pH environments generally exhibiting quite restrictive environments, where it may be hard for plants to establish and persist even under generally suitable conditions. For example, nitrogen becomes a limiting factor in more acidic soils causing low growth rates (e.g., Nordbakken et al. 2004; Pegtel 1994) and hence acididophilic communities could be more sensitive to human disturbance (for example drainage), possibly resulting in a higher plant dark diversity. Many plants adapted to low-pH soils such as carnivorous plants or heathland plants are adapted to natural disturbances (e.g., fire), and have poor establishment success when disturbances are absent or the competition from fast-growing plants is high (Brewer 1998; Nordbakken et al. 2004; Paniw et al. 2017; Pegtel 1994). For conservation projects in low-pH sites, this suggests that focus on natural processes facilitating successful establishment of characteristic plant species is central.

Our study showed that plant dark diversity was generally higher in sites with relatively tall vegetation and – in the open dry habitats – also in sites with higher soil nitrogen content and where the vegetation is homogenous. This is supported by recent studies, showing that plants preferring higher nutrient availability is also less often part of the dark diversity (Moeslund et al. 2017) and that plant dark diversity is generally higher in relatively nutrient rich sites (Fløjgaard et al. 2020). In nature, overgrowth by tall herbs is a severe threat to low-statured slowly growing plants as these are outcompeted through asymmetric competition (Freckleton and Watkinson 2001; Hautier et al. 2018; Lepš 1999). This could explain the aforementioned findings; high soil nitrogen may cause a tall monotonic vegetation outcompeting slow-growing species. Also, tall vegetation could indicate lack of natural grazing due to the lack of large herbivores in most of the European landscape (Fløjgaard et al. 2022). Alternatively, high soil nitrogen is symptomatic of former fertilization and hence probably ploughing which in combination has caused a change in vegetation towards taller vegetation and a more unnatural or stochastic species composition (e.g., Enríquez-de-Salamanca 2020; Harvolk et al. 2015; Muratet et al. 2008) causing a higher dark diversity.

In open landscapes, the plant dark diversity was often highest in sites with a relatively homogenous terrain (i.e., a high average terrain openness that only varies little). Homogenous terrains with gentle slopes are often more affected by human impacts (Odgaard et al. 2014; Sandel and Svenning 2013), and this could explain the relationship between homogenous terrains and dark diversity. Additionally, places with low variation in solar radiation input also often had a high dark diversity. This could reflect that niches related to heat load, light or soil moisture are important for local plant diversity (Moeslund et al. 2013a; Silvertown et al. 2015; Stein et al. 2014). Even small changes in these factors can have a relatively high impact on the vegetation (Moeslund et al. 2013a) and possibly the low solar radiation variation in high-dark diversity sites can reflect that small abiotic changes allow a number of species to thrive where they would otherwise not. On the other hand, high-dark diversity habitats also had relatively high local (0.5 m scale) terrain variations somewhat contrary to the common hypothesis that microtopography is related to the number of microhabitats and therefore important for local plant diversity (Moeslund et al. 2013b; Stein et al. 2014).

Tamme and colleagues (2010) proposed that that at fine spatial scales environmental heterogeneity can be negatively correlated with observed plant diversity possibly because microfragmentation effects restrains the establishment of new individuals. Likewise, such microfragmentation effects could explain the pattern we observed in this study; more species are missing when fine-scale topographical variation are high. Alternatively, this finding could reflect a higher number of unfilled microhabitats in sites where microtopographical variations are rather high.

#### Forest

Forests having relatively few natural-forest indicators (forest-indicator species, bryophyte species or number of rot attacks), also had a higher plant dark diversity. These natural-forest indicators probably reflect the degree to which natural processes are missing, e.g., natural hydrology (Maanavilja et al. 2014) or natural impacts by large herbivores (Kowalczyk et al. 2021). Hence, this relationship seems rather logic; the less natural, the more species are missing in the forest plots.

We found a higher dark diversity in tall forests. This could have at least two plausible explanations. Firstly, relatively tall and dense canopies are characteristic for production forests. So, this pattern could be due to forestry, altering the forest environment and thereby causing a higher dark diversity. Even though the included forests are in principle protected (NATURA2000), some are still being logged. Secondly, forests were not even-aged and this finding could simply reflect that taller forests are older and have a higher dark diversity, possibly due to a higher number of unfilled niches in these more developed woodlands (Lilja et al. 2006). However, this is against our hypothesis and seems counter-intuitive as species should have had time to establish in these older forests, so we consider the first explanation the most likely.

The dark diversity in forests had a non-linear relationship to the variation in vegetation cover, with the highest dark diversity found in forests with either high or low variation in this measure. If there is high variation, the number of unfilled niches may be quite high (Ashby et al. 2017) or the disturbances having caused these variations could have disturbed the flora to the extent where some species cannot cope with it, possibly due to microfragmentation effects (Tamme et al. 2010). Several recent studies found no effect of forest stand structural variation on local forest plant diversity (Penone et al. 2019; Zellweger et al. 2016), although in the light of the environmental heterogeneity hypothesis this would be expected (Stein et al. 2014). Perhaps this could also be explained by microfragmentation effects, and in that case together with our findings this suggests that such local scale fragmentation effects actually play important roles for forest plant diversity. In the second case, i.e. if there is consistently low variation in vegetation cover, fewer species tend to dominate the forest; for example in closed forests mostly trees will dominate (Thomas et al. 1999). In open forests with a low vegetation cover variation (i.e., mostly herb cover), it is likely that grasses or clonal tall herbs will dominate, as they will probably benefit from root gaps (Parsons et al. 1994; Thomas et al. 1999) or the decay of litter and tree stumps in old forest floors (Bohara et al. 2020; Prescott 2002). Indeed, this is likely in our study where 70.0 % of the plots (404 of 577) with closed forests and low variation in vegetation cover (plots < 25^th^ percentiles of canopy openness and vegetation cover variation) had few (< 2) species that forms dense monotonic vegetation (not shown, Table S2 in Supplementary Information lists these species). This was the case for only 29.8 % of the open forests (17 of 57 plots) with a low variation in vegetation cover (plots > 75^th^ percentile of canopy openness, same vegetation cover variation as above). Under natural conditions, open forests would be a great benefit for nature, and dark diversity would be expected to be low here. However, in Denmark, the impact from large herbivores is unnaturally low (Fløjgaard et al. 2021). Consequently, tall herbs forming monotonic stands often dominate, and many species that could otherwise thrive are missing.

### Field-vs. lidar-based measures

In this study, we showed that – while models including measured factors persistently performed better than lidar-only models – lidar provides important insight into the characteristics of high-dark-diversity habitats. This illustrates the strengths of lidar for characterizing habitats but also emphasizes that it cannot fully stand alone even when thoroughly using all the information in a lidar point cloud as we did here. This finding is supported by recent studies, e.g., by Valdez et al. (2021) who studied the characteristics of high-dark-diversity fungal communities and by Zellweger et al. (2016) who shed light on the relative importance of abiotic vs. structural factors for species richness at landscape scale in Swiss forests. It is notoriously difficult to estimate soil chemistry and -type using lidar (but see Li et al. 2016). We believe this is an important explanation for these findings. We foresee that future research and new possibilities both with point clouds and artificial intelligence and in other areas of remote sensing (i.e., fine spatial and spectral resolution data) could potentially bridge this gap.

## Acknowledgements

We sincerely thank Aage V. Jensen Nature Fund for financial support to JEM, CF, AKB, LD, and KC through the project “Dark Diversity in Nature Management”. MP was supported by the Estonian Research Council (PRG609) and the European Regional Development Fund (Centre of Excellence EcolChange).

## Data accessibility

The vegetation data used for this study is freely available for download through naturdata.miljoeportal.dk. Lidar data is freely available at datafordeler.dk.

## Notes

### Competing Interest Statement

The authors have declared no competing interest.

## Literature cited

Akaike, H. (1974). A new look at the statistical model identification. IEEE Transactions on Automatic Control, 19, 716–723

Ashby, B., Watkins, E., Lourenço, J., Gupta, S., & Foster, K.R. (2017). Competing species leave many potential niches unfilled. Nature Ecology & Evolution, 1, 1495–1501

Beals, E.W. (1984). Bray-Curtis ordination: an effective strategy for analysis of multivariate ecological data. Advances in Ecological Research, 14, 1–55

Blangiardo, M., Cameletti, M., Baio, G., & Rue, H. (2013). Spatial and spatio-temporal models with R-INLA. Spatial and Spatio-temporal Epidemiology, 7, 39–55

Bohara, M., Acharya, K., Perveen, S., Manevski, K., Hu, C., Yadav, R.K.P., Shrestha, K., & Li, X. (2020). In situ litter decomposition and nutrient release from forest trees along an elevation gradient in Central Himalaya. Catena, 194, 104698

Brewer, J.S. (1998). Effects of competition and litter on a carnivorous plant, Drosera capillaris (Droseraceae). American Journal of Botany, 85, 1592–1596

Brunbjerg, A.K., Bruun, H.H., Brøndum, L, Classen, A.T., Fog, K., Frøslev, T.G., Goldberg, I., Hansen, M.D.D., Høye, T.T., Læssøe, T., Newman, G., Skipper, L, Søchting, U., & Ejrnæs, R. (2019). A systematic survey of regional multitaxon biodiversity: evaluating strategies and coverage. BMC Ecology, 19, 43

Buchwald, E., Wind, P., Bruun, H.H., Møller, P.F., Ejrnæs, R., & Svart, H.E. (2013). Hvilke planter er hjemmehørende i Danmark? Flora & fauna, 118, 73–96

Burnham, K.P., & Anderson, D.R. (2002). Model Selection and Multimodel Inference: A Practical Information-Theoretic Approach. New York: Springer-Verlag

Council of the European Communities (1992). Council Directive 92/43/EEC of 21 May 1992 on the conservation of natural habitats and of wild fauna and flora. In Official Journal of the European Council, Series L (pp. 7–50)

de Vries, J.P.R., Koma, Z., WallisDeVries, M.F., & Kissling, W.D. (2021). Identifying fine-scale habitat preferences of threatened butterflies using airborne laser scanning. Diversity and Distributions, 27, 1251–1264

Depauw, L, Perring, M.P., Brunet, J., Maes, S.L., Blondeel, H., De Lombaerde, E., De Groote, R., & Verheyen, K. (2019). Interactive effects of past land use and recent forest management on the understorey community in temperate oak forests in South Sweden. Journal of Vegetation Science, 30, 917–928

Doneus, M. (2013). Openness as visualization technique for interpretative mapping of airborne lidar derived digital terrain models. Remote Sensing, 5, 6427–6442

Enríquez-de-Salamanca, Á. (2020). Human influence on the flora of the Spanish Central Range. Plant Biosystems-An International Journal Dealing with all Aspects of Plant Biology, 154, 474–480

Fløjgaard, C., Buttenschøn, R.M., Byriel, D.B., Clausen, K.K., Gottlieb, L, Kanstrup, N., Strandberg, B., & Ejrnæs, R. (2021). Biodiversitetseffekter af rewilding. Videnskabelig rapport nr. 425, 124 pp.

Fløjgaard, C., Pedersen, P.B.M., Sandom, C.J., Svenning, J.-C., & Ejrnæs, R. (2022). Exploring a natural baseline for large-herbivore biomass in ecological restoration. Journal of Applied Ecology, 59, 18–24

Fløjgaard, C., Valdez, J., Dalby, L., Moeslund, J.E., Clausen, K., Ejrnæs, R., Pärtel, M., & Brunbjerg, A.K. (2020). Dark diversity reveals importance of biotic resources and competition for plant diversity across habitats. Ecology and Evolution, 10, 6078–6088

Freckleton, R.P., & Watkinson, A.R. (2001). Asymmetric competition between plant species. Functional Ecology, 15, 615–623

Gijbels, P., Adriaens, D., & Honnay, O. (2012). An Orchid Colonization Credit in Restored Calcareous Grasslands. Ecoscience, 19, 21–28

Hanson, J.O., Rhodes, J.R., Butchart, S.H.M., Buchanan, G.M., Rondinini, C., Ficetola, G.F., & Fuller, R.A. (2020). Global conservation of species’ niches. Nature, 580, 232–234

Harvolk, S., Symmank, L, Sundermeier, A., Otte, A., & Donath, T.W. (2015). Human impact on plant biodiversity in functional floodplains of heavily modified rivers–A comparative study along German Federal Waterways. Ecological Engineering, 84, 463–475

Hautier, Y., Vojtech, E., & Hector, A. (2018). The importance of competition for light depends on productivity and disturbance. Ecology and Evolution, 8, 10655–10661

Höfle, B., Hollaus, M., & Hagenauer, J. (2012). Urban vegetation detection using radiometrically calibrated small-footprint full-waveform airborne LiDAR data. ISPRS Journal of photogrammetry and remote sensing, 67, 134–147

Junttila, S., Sugano, J., Vastaranta, M., Linnakoski, R., Kaartinen, H., Kukko, A., Holopainen, M., Hyyppä, H., & Hyyppä, J. (2018). Can leaf water content be estimated using multispectral terrestrial laser scanning? A case study with Norway spruce seedlings. Frontiers in plant science, 9, 299

Kowalczyk, R., Kamiński, T., & Borowik, T. (2021). Do large herbivores maintain open habitats in temperate forests? Forest Ecology and Management, 494, 119310

Lepš, J. (1999). Nutrient status, disturbance and competition: an experimental test of relationships in a wet meadow. Journal of Vegetation Science, 10, 219–230

Lewis, R.J., de Bello, F., Bennett, J.A., Fibich, P., Finerty, G.E., Götzenberger, L., Hiiesalu, I., Kasari, L., Lepš, J., Májeková, M., Mudrák, O., Riibak, K., Ronk, A., Rychtecká, T., Vitová, A., & Pärtel, M. (2017). Applying the dark diversity concept to nature conservation. Conservation Biology, 31, 40–47

Lewis, R.J., Szava-Kovats, R., & Pärtel, M. (2015). Estimating dark diversity and species pools: an empirical assessment of two methods. Methods in Ecology and Evolution, 7, 104–113

Li, C., Xu, Y., Liu, Z., Tao, S., Li, F., & Fang, J. (2016). Estimation of Forest Topsoil Properties Using Airborne LiDAR-Derived Intensity and Topographic Factors. Remote Sensing, 8, 561

Lilja, S., Wallenius, T., & Kuuluvainen, T. (2006). Structure and development of old Picea abies forests in northern boreal Fennoscandia. Ecoscience, 13, 181–192

Lindgren, F., Rue, H., & Lindström, J. (2011). An explicit link between Gaussian fields and Gaussian Markov random fields: the stochastic partial differential equation approach. Journal of the Royal Statistical Society: Series B (Statistical Methodology), 73, 423–498

McCune, B., & Keon, D. (2002). Equations for potential annual direct incident radiation and heat load. Journal of Vegetation Science, 13, 603–606

Menard, S. (2001). Applied Logistic Regression Analysis. (2nd edition ed.). SAGE Publications, Inc

Moeslund, J.E., Arge, L, Bøcher, P.K., Dalgaard, T., Odgaard, M.V., Nygaard, B., & Svenning, J.-C. (2013a). Topographically controlled soil moisture is the primary driver of local vegetation patterns across a lowland region. Ecosphere, 4, Article 91

Moeslund, J.E., Arge, L, Bøcher, P.K., Dalgaard, T., & Svenning, J.-C. (2013b). Topography as a driver of local terrestrial vascular plant diversity patterns. Nordic Journal of Botany, 31, 129–144

Moeslund, J.E., Brunbjerg, A.K., Clausen, K.K., Dalby, L, Fløjgaard, C., Juel, A., & Lenoir, J. (2017). Using dark diversity and plant characteristics to guide conservation and restoration. Journal of Applied Ecology, 54, 1730–1741

Moeslund, J.E., Zlinszky, A., Ejrnæs, R., Brunbjerg, A.K., Bøcher, P.K., Svenning, J.-C., & Normand, S. (2019). Light detection and ranging explains diversity of plants, fungi, lichens, and bryophytes across multiple habitats and large geographic extent. Ecological Applications, 29, e01907

Morel, L., Jung, V., Chollet, S., Ysnel, F., & Barbe, L. (2022). From taxonomic to functional dark diversity: Exploring the causes of potential biodiversity and its implications for conservation. Journal of Applied Ecology, 59, 103–116

Muratet, A., Porcher, E., Devictor, V., Arnal, G., Moret, J., Wright, S., & Machon, N. (2008). Evaluation of floristic diversity in urban areas as a basis for habitat management. Applied Vegetation Science, 11, 451–460

Münzbergová, Z., & Herben, T. (2004). Identification of suitable unoccupied habitats in metapopulation studies using co-occurrence of species. Oikos, 105, 408–414

Mäyrä, J., Keski-Saari, S., Kivinen, S., Tanhuanpää, T., Hurskainen, P., Kullberg, P., Poikolainen, L., Viinikka, A., Tuominen, S., Kumpula, T., & Vihervaara, P. (2021). Tree species classification from airborne hyperspectral and LiDAR data using 3D convolutional neural networks. Remote Sensing of Environment, 256, 112322

Maanavilja, L., Aapala, K., Haapalehto, T., Kotiaho, J.S., & Tuittila, E.-S. (2014). Impact of drainage and hydrological restoration on vegetation structure in boreal spruce swamp forests. Forest Ecology and Management, 330, 115–125

Nordbakken, J.-F., Rydgren, K., & Økland, R.H. (2004). Demography and population dynamics of Drosera anglica and D. rotundifolia. Journal of Ecology, 92, 110–121

Odgaard, M.V., Bøcher, P.K., Dalgaard, T., Moeslund, J.E., & Svenning, J.-C. (2014). Human-driven topographic effects on the distribution of forest in a flat, lowland agricultural region. Journal of Geographical Sciences, 24, 76–92

Oksanen, J., Blanchet, F.G., Friendly, M., Kindt, R., Legendre, P., McGlinn, D., Minchin, P.R., O’Hara, R.B., Simpson, G.L., Solymos, P., Stevens, M.H.H., Szoecs, E., & Wagner, H. (2017). vegan: Community Ecology Package. R package version 2.4–3

Paniw, M., Quintana-Ascencio, P.F., Ojeda, F., & Salguero-Gómez, R. (2017). Interacting livestock and fire may both threaten and increase viability of a fire-adapted Mediterranean carnivorous plant. Journal of Applied Ecology, 54, 1884–1894

Paradis, E., & Schliep, K. (2018). ape 5.0: an environment for modern phylogenetics and evolutionary analyses in R. Bioinformatics, 35, 526–528

Parsons, W.F.J., Knight, D.H., & Miller, S.L. (1994). Root Gap Dynamics in Lodgepole Pine Forest: Nitrogen Transformations in Gaps of Different Size. Ecological Applications, 4, 354–362

Pegtel, D.M. (1994). Habitat characteristics and the effect of various nutrient solutions on growth and mineral nutrition of *Arnica montana* L. grown on natural soil. Vegetatio, 114, 109–121

Penone, C., Allan, E., Soliveres, S., Felipe-Lucia, M.R., Gossner, M.M., Seibold, S., Simons, N.K., Schall, P., van der Plas, F., Manning, P., Manzanedo, R.D., Boch, S., Prati, D., Ammer, C., Bauhus, J., Buscot, F., Ehbrecht, M., Goldmann, K., Jung, K., Müller, J., Müller, J.C., Pena, R., Polle, A., Renner, S.C., Ruess, L., Schönig, I., Schrumpf, M., Solly, E.F., Tschapka, M., Weisser, W.W., Wubet, T., & Fischer, M. (2019). Specialisation and diversity of multiple trophic groups are promoted by different forest features. Ecology Letters, 22, 170–180

Pfeifer, N., Mandlburger, G., Otepka, J., & Karel, W. (2014). OPALS – A framework for Airborne Laser Scanning data analysis. Computers, Environment and Urban Systems, 45, 125–136

Prescott, C.E. (2002). The influence of the forest canopy on nutrient cycling. Tree Physiology, 22, 1193–1200

Príncipe, A., Matos, P., Sarris, D., Gaiola, G., do Rosário, L., Correia, O., & Branquinho, C. (2019). In Mediterranean drylands microclimate affects more tree seedlings than adult trees. Ecological Indicators, 106, 105476

Pärtel, M. (2002). Local plant diversity patterns and evolutionary history at the regional scale. Ecology, 83, 2361–2366

Pärtel, M., Szava-Kovats, R., & Zobel, M. (2011). Dark diversity: shedding light on absent species. Trends in Ecology & Evolution, 26, 124–128

Ratajczak, Z., Nippert, J.B., & Collins, S.L. (2012). Woody encroachment decreases diversity across North American grasslands and savannas. Ecology, 93, 697–703

Riibak, K., Reitalu, T., Tamme, R., Helm, A., Gerhold, P., Znamenskiy, S., Bengtsson, K., Rosén, E., Prentice, H.C., & Pärtel, M. (2015). Dark diversity in dry calcareous grasslands is determined by dispersal ability and stress-tolerance. Ecography, 38, 713–721

Ronk, A., Szava-Kovats, R., & Pärtel, M. (2015). Applying the dark diversity concept to plants at the European scale. Ecography, 38, 1015–1025

Rue, H., Martino, S., & Chopin, N. (2009). Approximate Bayesian inference for latent Gaussian models by using integrated nested Laplace approximations. Journal of the Royal Statistical Society: Series B (Statistical Methodology), 71, 319–392

Sandel, B., & Svenning, J.-C. (2013). Human impacts drive a global topographic signature in tree cover. Nature Communications, 4, 2474

Silvertown, J., Araya, Y., & Gowing, D. (2015). Hydrological niches in terrestrial plant communities: a review. Journal of Ecology, 103, 93–108

Stein, A., Gerstner, K., & Kreft, H. (2014). Environmental heterogeneity as a universal driver of species richness across taxa, biomes and spatial scales. Ecology Letters, 17, 866–880

Tamme, R., Hiiesalu, L, Laanisto, L., Szava-Kovats, R., & Pärtel, M. (2010). Environmental heterogeneity, species diversity and co-existence at different spatial scales. Journal of Vegetation Science, 21, 796–801

Team, R.C. (2020). R: A language and environment for statistical computing. R foundation for Statistical Computing, Vienna, Austria

Thomas, S.C., Halpern, C.B., Falk, D.A., Liguori, D.A., & Austin, K.A. (1999). Plant diversity in managed forests: understory responses to thinning and fertilization. Ecological Applications, 9, 864–879

Valdez, J.W., Brunbjerg, A.K., Fløjgaard, C., Dalby, L, Clausen, K.K., Pärtel, M., Pfeifer, N., Hollaus, M., Wimmer, M.H., Ejrnæs, R., & Moeslund, J.E. (2021). Relationships between macro-fungal dark diversity and habitat parameters using LiDAR. Fungal Ecology, 51, 101054

Wen, Z., Cai, T., Feijó, A., Xia, L., Cheng, J., Ge, D., & Yang, Q. (2020). Using completeness and defaunation indices to understand nature reserve’s key attributes in preserving medium-and large-bodied mammals. Biological Conservation, 241, 108273

Zellweger, F., Baltensweiler, A., Ginzler, C., Roth, T., Braunisch, V., Bugmann, H., & Bollmann, K. (2016). Environmental predictors of species richness in forest landscapes: abiotic factors versus vegetation structure. Journal of Biogeography, 43, 1080–1090

Zlinszky, A., Schroiff, A., Kania, A., Deák, B., Mücke, W., Vári, Á., Székely, B., & Pfeifer, N. (2014). Categorizing grassland vegetation with full-waveform airborne laser scanning: A feasibility study for detecting Natura 2000 habitat types. Remote Sensing, 6, 8056–8087

Zobel, M. (2016). The species pool concept as a framework for studying patterns of plant diversity. Journal of Vegetation Science, 27, 8–18

Zuur, A.F., Ieno, E.N., & Seveliev, A.A. (2017). Beginner’s guide to spatial, temporal and spatial-temporal ecological data analysis with R-INLA. Newburgh, UK: Highland Statistics

