## Supporting Information for "The characteristics of high-dark-diversity habitats derived from lidar"

***Supporting information for the paper entitled "The characteristics of high-dark-diversity habitats derived from lidar"***

**Description of how field-measured variables were obtained in the dataset used for this study**

**pH**: For most dry open habitats the pH was measured from a soil sample in a certified lab. Soil samples were taken at the four corners of a central 1x1 m quadrat in each plot in the upper 5 cm of soil. Before soil sampling all loose litter, twigs, leaves etc. was removed. The four soil samples were subsequently blended, air-dried, stored cool and sent to a certified lab for standard pH analysis. For wet habitats the pH was measured directly in the field from a water sample using a calibrated pH meter. Water was sampled using a piezometer tube that was hammered into the root zone of the soil. After water entered into the tube, it was sampled into a container using a hose. The first water was used for rinsing the contained and then discarded. Pollution with surface-water was prevented to the degree possible, so for example sampling in flooded areas was not conducted. The water pH measurements have an estimated uncertainty of 15 %.

**Nitrogen**: For most dry open habitats, nitrogen was measured in a soil sample in a certified lab. Part of the same soil sample as used for pH measurement was used for this analysis, which was a standard total nitrogen content analysis with an uncertainty of 20 %. For the some of the wet habitats, nitrogen was estimated by measuring nitrate-N in water samples. The water samples for this were taken in a manner similar to that for pH, but the water was filtered through a filter approved by the lab to where samples was handed in subsequently, for example CAMEO 30GA. Water was stored dark and between 0 and 4 degrees celsius and handed in to the lab within 24 hours. The lab conducted a standard nitrate-N analysis with 15 % uncertainty.

**Soil carbon**: Soil carbon was measured using a part of the same soil sample as used for pH and soil nitrogen measurements. The lab conducted a standard total carbon content analysis with an uncertainty of 15 %.

**Cover of woody vegetation below and above 1 m**: The cover of woody plants below and above 1 m respectively were visually estimated. All species belonging to *Rubus* subgenus *Rubus* (blackberry) and *Rubus ideaus* did not count as woody species.

**Number of forest indicator species**: The following twenty-five fungi, lichen and bryophyte species count as forest indicator species and was counted for each forest plot: *Daedaleopsis confragosa* (Bolton) J. Schröt., *Eutypa spinosa* (Pers.) Tul. & C. Tul., *Fomes fomentarius* (L.) J.J. Kickx, *Fomitopsis pinicola* (Sw.) P. Karst., *Ganoderma applanatum* (Pers.) Pat., *Ganoderma pfeifferi* Bres., *Hymenochaete rubiginosa* (Dicks.) Lév., *Mensularia radiata* (Sowerby) Lázaro Ibiza, *Inocutis rheades* (Pers.) Fiasson & Niemelä, *Ischnoderma resinosum* (Schrad.) P. Karst., *Phellinus tremulae* (Bondartsev) Bondartsev & Borissov, *Piptoporus betulinus* (Bull.) P. Karst., *Lecanactis abietina* (Ach.) Körb., *Lobaria pulmonaria* (L.) Hoffm., *Opegrapha vermicellifera* (Kunze) J.R. Laundon, *Pyrenula nitida* (Weigel) Ach., *Thelotrema lepadinum* (Ach.) Ach., *Homalothecium sericeum* (Hedw.) Schimp., *Isothecium alopecuroides* (Lam. ex Dubois) Isov., *Isothecium myosuroides* Brid., *Neckera complanata* (Hedw.) Huebener, *Plagiochila asplenioides* (L. emend. Taylor) Dumort. subsp. *asplenioides*, *Porella platyphylla* (L.) Pfeiff., *Rhytidiadelphus loreus* (Hedw.) Warnst., *Zygodon* species.

**Number of hollows and rot attacks**: Within each forest plot the number of living trees with hollows and rotten areas was recorded. The same trees can have both hollows and rotten areas and was hence counted both for number of hollows and number of rot attacks. Hollows and rotten areas were only counted when they occured on the tree main stem or on branches with a diameter above 20 cm, and only those occurring above 0.5 m above ground and visible from the ground without using a ladder were counted. A hollow was defined as a hole in the bark with underlying rot attack or a hollow to a depth of at least 5 cm. Rotten areas were defined as areas above 100 cm2 with loose bark or naked wood that is clearly decaying. Dead branches can leave such areas and they were counted if the branch diameter was above 11 cm. Fresh bark wounds did not count.

**Supporting tables**

**Table S1.** Measures kept in model selection (X), removed in Variance Inflation Factor analysis (-), removed since we had no hypothesis for including them (*) or deselected in subsequent model selection (/)

| | pH | Nitrogen | Soil carbon | Cover of woody vegetation < 1 m | Cover of woody vegetation > 1 m | No. of forest indicator species | No. of bryophyte species | No. of hollows | No. of rot attacks | Terrain roughness | Terrain slope | Terrain slope var. | Terrain openness | Terrain openness var. | Potential solar radiation | Potential solar radiation var. | Heat load | Heat load var. | Vegetation height | Vegetation height var. | Vegetation cover | Vegetation cover var. | Echo ratio | Echo ratio var. | Canopy openness | Canopy openness var. | Lidar amplitude |
|---|---|---|---|---|---|---|---|---|---|---|---|---|---|---|---|---|---|---|---|---|---|---|---|---|---|---|---|
| Open dry F + L | X | X | - | | | | | | | X | - | - | X | / | / | X | / | X | - | - | / | - | - | - | - | X | / |
| Open dry L | | | | | | | | | | X | - | - | X | X | X | X | X | X | X | X | - | - | - | - | - | - | / |
| Open wet F + L | X | / | / | | | | | | | X | - | - | X | X | X | X | / | X | X | - | - | - | - | - | X | / | / |
| Open wet L | | | | | | | | | | X | - | - | X | X | X | X | X | X | X | X | - | - | - | - | - | - | / |
| Open all L | | | | | | | | | | X | - | - | X | X | X | X | X | X | X | X | - | - | - | - | - | - | / |
| Forest reduced F + L | | | | X | X | X | X | / | X | / | - | X | * | * | X | - | X | / | X | X | / | X | - | - | / | - | X |
| Forest reduced L | | | | | | | | | | / | - | / | * | * | - | - | / | / | X | X | / | / | - | - | X | X | X |
| Forest all L | | | | | | | | | | / | - | / | * | * | - | - | / | / | X | X | / | / | - | - | X | X | X |
| Transformation (open models, dry) | Ac | Inv | Ln | | | | | | | No | No | No | No | No | No | Ln | No | No | No | No | No | Inv | No | No | No | No | No |
| Transformation (open models, dry) | Ac | No | Ln | | | | | | | No | No | No | No | No | No | Ln | No | No | No | No | No | Inv | No | No | No | No | No |
| Transformation (forest models) | | | | Ln | No | Ln | Ln | Ln | Ln | Sq | Ac | No | No | Ln | No | No | No | No | No | Sq | No | No | Ln | No | No | No | No |

F: Field-based measures, L: lidar-based measures. For example, Open dry M + L means the model for the open landscapes with primarily dry habitat types and considering both field- and lidar-based measures. Ac: arc-cosinus, Inv: inverse, No: no transformation, Ln: log, Sq: square root.

**Table S2.** List of species found in the forest plots of this study and that are known to form vegetation patches (dense monotonic vegetation). Growth description is based on Mossberg and Stenberg (2020). Asterisks mark species where the growth description was not taken from the above reference but described from the authors' field experience.

| **Species name** | **Growth** |
|---|---|
| *Aegopodium podagraria* L. | Forms dense monospecific carpets |
| *Agrostis canina* L. | Loose tussocks, extensive stolons |
| *Agrostis capillaris* L. | Loose tussocks or loose monospecific vegetation |
| *Agrostis stolonifera* L. | Extensive stolons |
| *Allium ursinum* L. | Forms dense monospecific carpets |
| *Alopecurus geniculatus* L. | Extensive stolons |
| *Anemone nemorosa* L. | Forms dense monospecific carpets |
| *Avenella flexuosa* (L.) Drejer | Loose tussocks, often dominating |
| *Calamagrostis canescens* (Weber) Roth | Forms dense monospecific carpets |
| *Calamagrostis epigejos* (L.) Roth | Forms open carpets |
| *Calluna vulgaris* (L.) Hull | Often dominating on poor sandy soil* |
| *Chamaenerion angustifolium* (L.) Scop. | Often forms monospecific stands* |
| *Cytisus scoparius* (L.) Link | Often forms monospecific stands* |
| *Elytrigia repens* (L.) Desv. ex Nevski | Forms dense monospecific carpets |
| *Festuca rubra* L. | Forms dense monospecific carpets |
| *Holcus mollis* L. | Forms dense monospecific carpets |
| *Lolium perenne* L. | Open tussocks or loose carpets |
| *Molinia caerulea* (L.) Moench | Dense tussocks often in carpets |
| *Petasites albus* (L.) Gaertn. | Forms dense monospecific carpets |
| *Petasites hybridus* (L.) G. Gaertn., B. Mey. & Scherb. | Forms dense monospecific carpets |
| *Poa pratensis* L. | Monospecific carpets, extensive stolons |
| *Pteridium aquilinum* (L.) Kuhn | Forms monospecific stands* |

### Supporting figures

**Figure S3.** Overview of all the open habitats covered in this study with sample size (n) and their regional pool adjusted dark diversity.

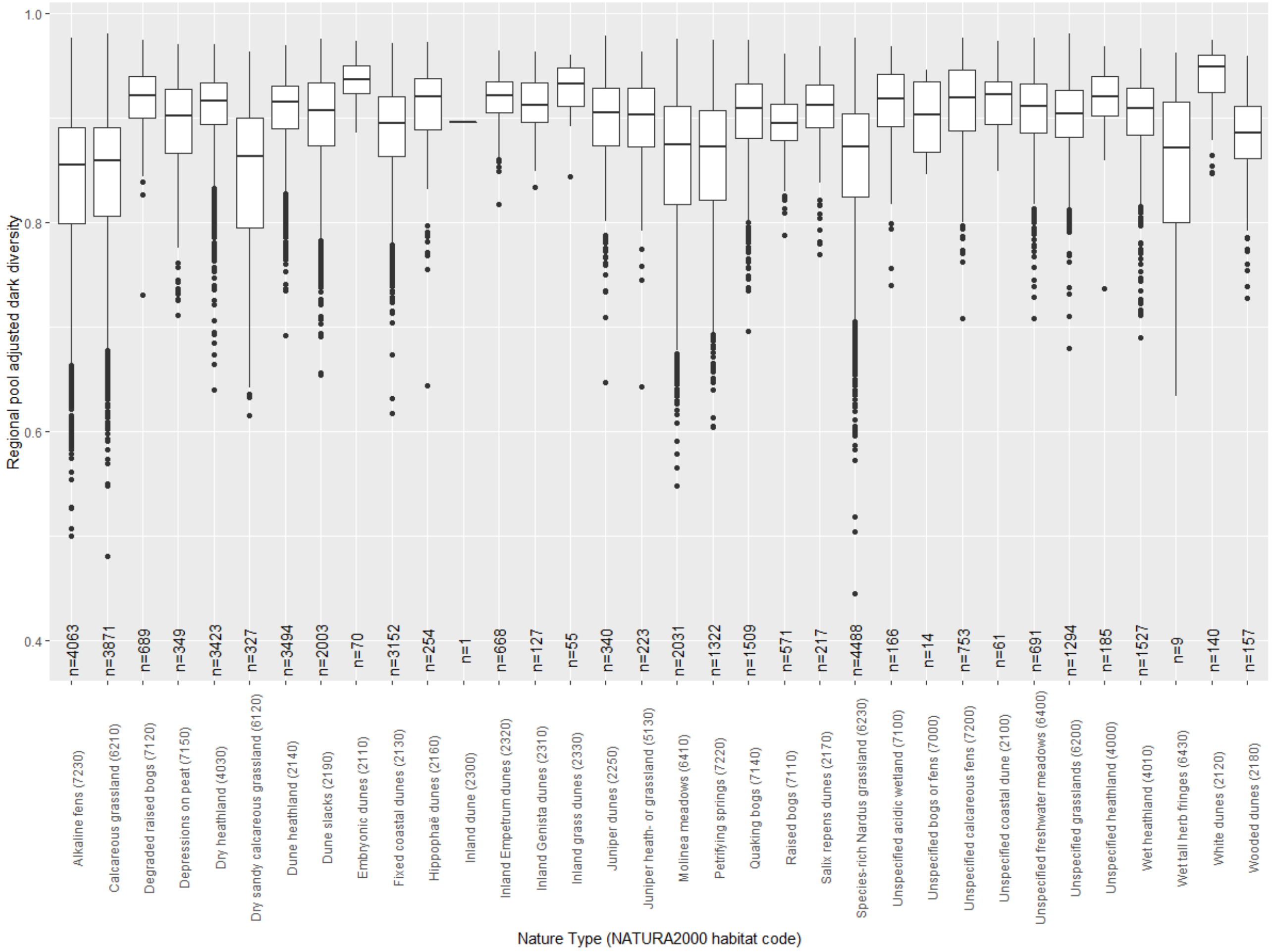

**Figure S4.** Overview of all the forest habitats covered in this study with sample size (n) and their regional pool adjusted dark diversity.

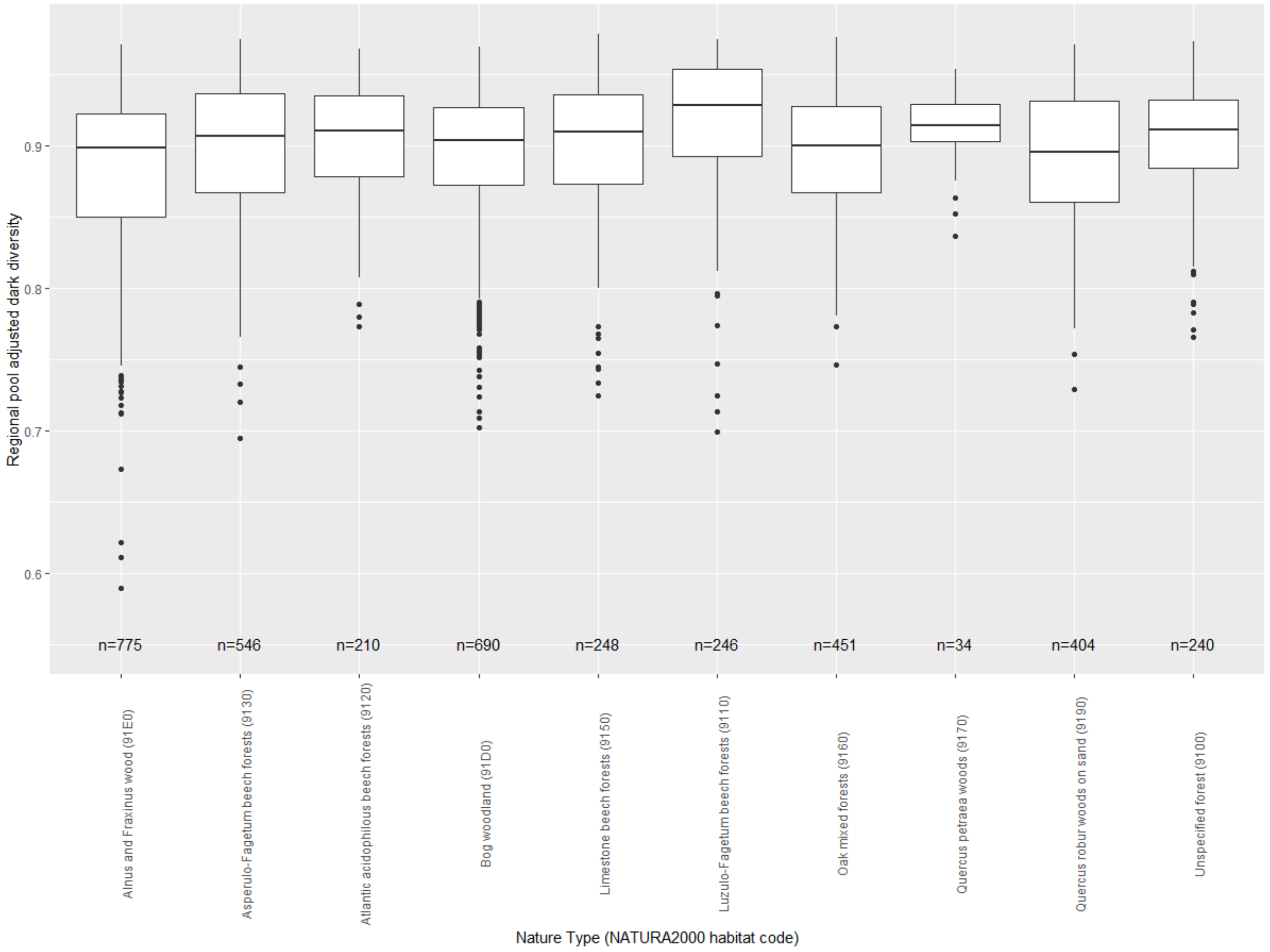

**Figure S5.** The derived range for spatial autocorrelation for each integrated nested Laplace approximation model for open landscapes

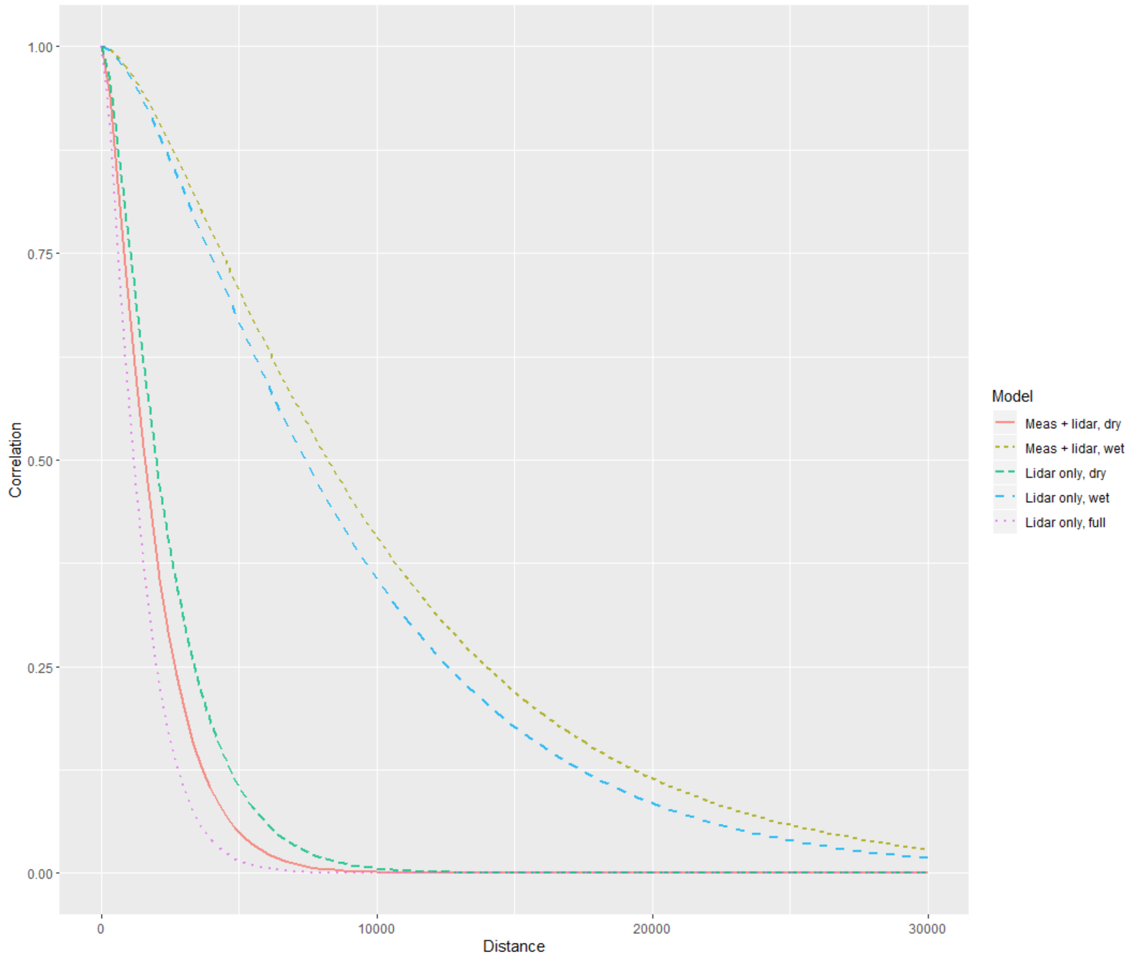

**Figure S6.** The derived range for spatial autocorrelation for each integrated nested Laplace approximation model for forests

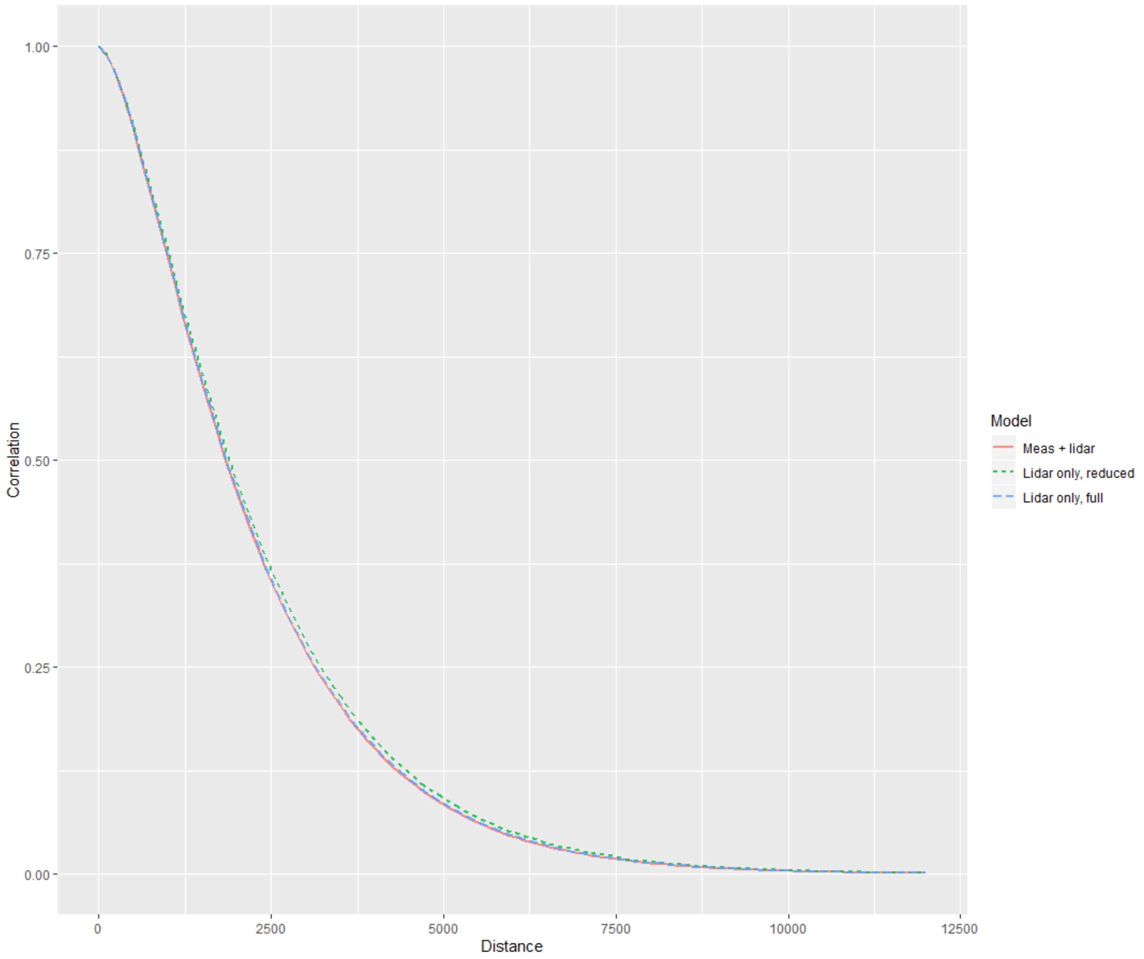

### Literature cited

Mossberg, B., & Stenberg, L. (2020). *Nordens flora*. (1st ed. ed.). Denmark: Gyldendal
